# Genetic timestamping of plasma cells *in vivo* reveals homeostatic population turnover

**DOI:** 10.1101/2020.04.12.038380

**Authors:** AQ Xu, RR Barbosa, DP Calado

## Abstract

Plasma cells (PC)s are essential for protection from infection, and at the origin of incurable cancers. Current studies do not circumvent limitations of removing PCs from their microenvironment and confound formation and maintenance. This is in part due to the lack of tools to perform specific genetic manipulation *in vivo*. Also, studies of PC population dynamics have mostly relied on the use of nucleotide analog incorporation that does not label quiescent cells, a property of most PCs. Here we characterize in detail a genetic tool (*Jchain*^creERT2^) that permits first-ever specific genetic manipulation in PC *in vivo*, across immunoglobulin isotypes. Using this tool we found that PC numbers remained constant over-time and that PC decay was compensated by the emergence of new cells, supporting an homeostatic turnover of the population. The *Jchain*^creERT2^ genetic tool paves the way for in-depth mechanistic understanding of PC biology and pathology *in vivo*, in their microenvironment.

**Highlights:** *Jchain* expression occurs in most plasma cells across immunoglobulin isotypes

*Jchain*^creERT2^ mediated genetic manipulation is effective only in plasma cells

Genetic timestamping of plasma cells reveals homeostatic regulation

## Introduction

Antibodies produced by plasma cells (PCs) are crucial for immune protection against infection and for vaccination success (Nutt et al., 2015). Upon activation B cells terminally differentiate into PCs, a process initiated by the downregulation of the B cell transcription factor PAX5 (Kallies et al., 2007). This event allows the expression of multiple factors normally repressed by PAX5, including *Xbp1*and *Jchain* (Castro and Flajnik, 2014; Nutt et al., 2015; Rinkenberger et al., 1996; Shaffer et al., 2004). PAX5 downregulation is also followed by the expression of the transcription factors IRF4 and BLIMP1 that play essential roles in the establishment of the PC program (Kallies et al., 2007; Klein et al., 2006; Shapiro-Shelef et al., 2003). Beyond physiology, multiple cancers have a PC as cell of origin, including multiple myeloma, the second most frequent hematological malignancy overall, for which a cure remains to be found (Palumbo and Anderson, 2011). As consequence, the study of gene function in PC biology and pathology is a subject of intense investigation.

However, at least in part because of technical limitations most PC studies make use of *in vitro* and cell transfer systems that remove PCs from their microenvironment. Currently, genetic manipulation of PCs is not specific and targets other cell populations such as B cells, confounding PC formation and maintenance. Also, studies on the global turnover of the PC population are lacking, as investigation of the regulation of PC maintenance has mostly relied on the use of nucleotide analogs that do not track the vast majority of PCs due to their quiescent nature.

We found that amongst well-known PC associated genes, *Jchain* (*Igj*) had the highest level and most specific expression in PC populations. JCHAIN is a small polypeptide required to multimerize IgM and IgA, and necessary for the transport of these Ig classes across the mucosal epithelium in a poly-Ig receptor mediated process (Brandtzaeg and Prydz, 1984; Castro and Flajnik, 2014; Max and Korsmeyer, 1985). Here we characterized in detail a GFP-tagged creERT2 allele at the *Jchain* endogenous locus: *Igj*^*creERT2*^, hereafter termed *Jchain*^*creERT2*^. We found at the single cell level that *Jchain* expression occurred in PCs across immunoglobulin isotypes, including IgG1. Using the *Jchain*^*creERT2*^ allele we performed the first-ever highly specific cre-loxP genetic manipulation in PCs residing in their natural microenvironment *in vivo*. Using this system, that allows inclusive genetic timestamping of plasma cells, independent of their cell cycle status, we uncovered that the numbers of PCs in the spleen and bone marrow remained constant over-time and that PC decay was compensated by the emergence of new PCs. These data suggest that the turnover of the plasma cell population is homeostatically regulated. The *Jchain*^creERT2^ is thus a validated plasma cell specific genetic tool that paves the way for in-depth mechanistic understanding of PC biology and pathology *in vivo*, in their microenvironment.

## Results

### *Jchain* transcripts are highly enriched in plasma cells

B-to-PC differentiation is a process that involves a complex network of factors (**Fig. 1A**; (Nutt et al., 2015)). We investigated the level and specificity of the expression of genes associated with PCs (*Xbp1, Jchain, Scd1, Irf4*, and *Prdm1*) through the analysis of a publicly available RNA sequencing dataset for immune cell populations (ImmGen, (Heng et al., 2008)). We first determined the cell populations with the highest transcript level for each factor. *Xbp1* and *Irf4* were primarily expressed in PCs, however the expression in bone marrow PCs (B_PC_BM) was less than 2-fold greater than that of non-PC populations (**Fig. 1B**). The expression of *Sdc1* and *Prdm1* was not specific to PCs (**Fig. 1C**). Notably, peritoneal cavity macrophages (MF_226+II+480lo_PC) expressed more *Sdc1* than bone marrow PCs (B_PC_BM), and a subset of FOXP3^+^ T cells (Treg_4_FP3+_Nrplo_Co) expressed higher levels of *Prdm1* than that observed in splenic plasmablasts (B_PB_Sp) and bone marrow PCs (B_PC_BM; **Fig. 1C**). In contrast, *Jchain* had the highest level of transcript expression in PCs compared to non-PCs and was the most PC specific amongst all factors, with a 40-fold enrichment over germinal center (GC) B cells (B_GC_CB_Sp; **Fig. 1D**). We concluded that the *Jchain* locus was a suitable candidate for the generation of PC specific genetic tools.

**Figure 1.**
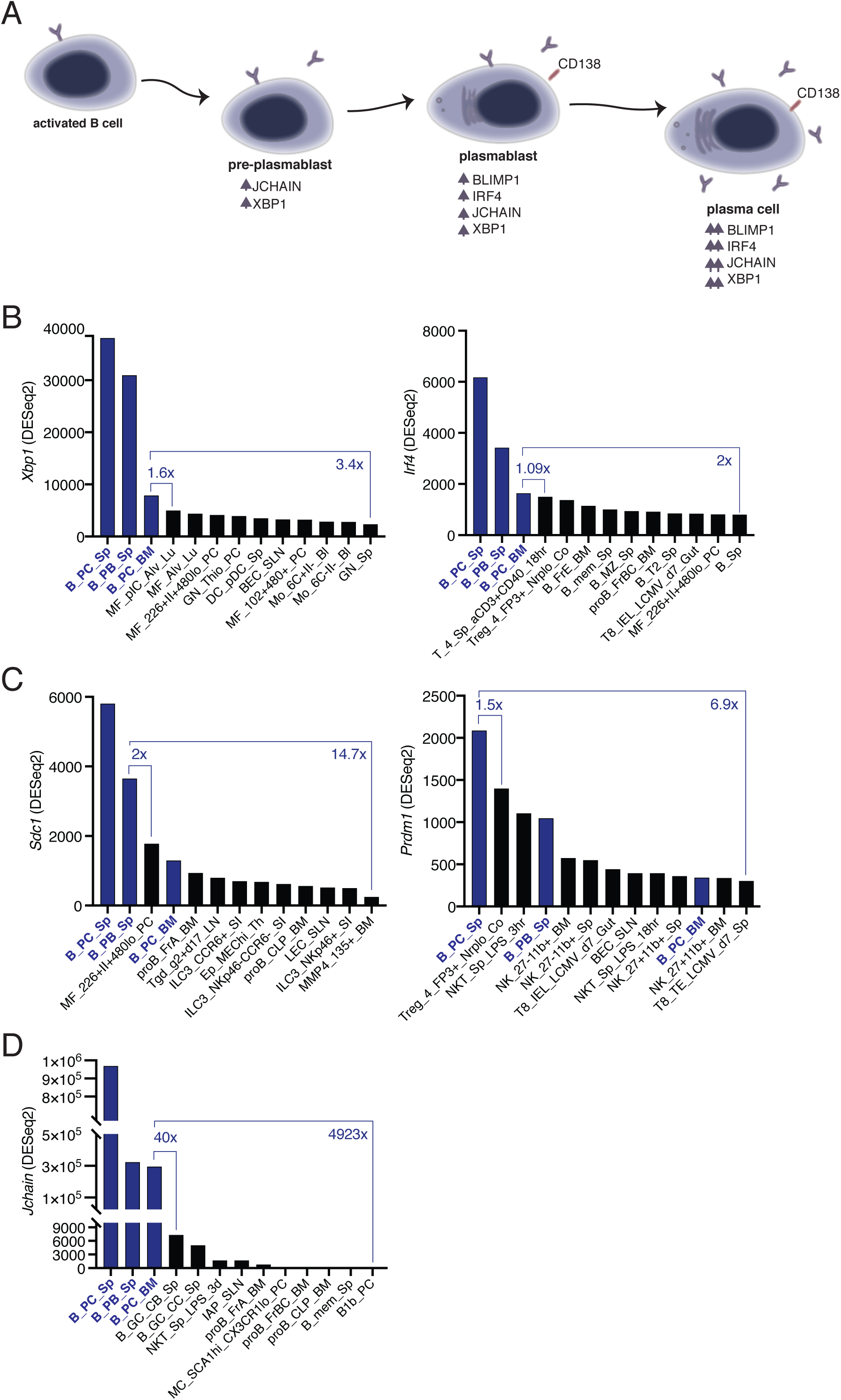
*Jchain* transcripts are highly enriched in plasma cells. **(A)** Schematic of the network of factors associated with plasma cell differentiation. Upwards arrows indicate increased expression compared to the precursor population. **(B-D)** Differential gene expression analysis using RNA sequencing data (ImmGen, (Heng et al., 2008)). (B) Analysis of *Xbp1* and *Irf4*, which encode for XBP1 and IRF4, respectively. (C) Analysis of *Sdc1* and *Prdm1*, which encode for CD138 and BLIMP1, respectively. (D) Analysis for *Jchain* (*Igj*) that encodes for JCHAIN. “x” indicates fold change. Expression Value Normalized by DESeq2. http://rstats.immgen.org/Skyline/skyline.html.

### *Jchain* is expressed in a small fraction of GC B cells and in most plasma cells

We searched alleles produced by the EUCOMMTools consortium and identified a genetically engineered *Jchain* allele produced by the Wellcome Trust Sanger Institute: MGI:5633773, hereafter termed *Jchain*^creERT2^. The genetically engineered *Jchain* allele contained an *FRT* site between exons 1 and 2 followed by an engrailed 2 splice acceptor sequence and an *EGFP*.*2A*.*cre*^*ERT2*^ expression cassette (**Fig. 2A**). In this design the expression of the EGFP (GFP) and of creERT2 is linked by a self-cleaving 2A peptide under the transcriptional control of the *Jchain* promoter (**Fig. 2A**). To determine cells with GFP expression we initially analyzed the spleen from mice heterozygous for the *Jchain*^creERT2^ allele that had been immunized with sheep red blood cells (SRBC) 12 days earlier **(Fig. 2B)**. B220 is expressed on the surface of B cells and downregulated during PC differentiation (Pracht et al., 2017). We therefore defined three cell populations based on the levels of GFP fluorescence and B220 surface expression: GFP^low^B220^high^, GFP^int^B220^int^, GFP^high^B220^low^, and a population negative for both markers (GFP^neg^B220^neg^; **Fig. 2C**). We next determined the fraction of cells within these populations That expressed surface CD138, a commonly used marker to define PCs by flow-cytometry (Pracht et al., 2017). The GFP^neg^B220^neg^ population did not contain CD138^+^ cells, however, the fraction of CD138^+^ cells increased in the remaining populations in agreement with the reduction of B220 expression during PC differentiation, and the GFP^int^B220^int^ and GFP^high^B220^low^ populations were mostly composed of CD138^+^ cells (**Fig. 2D** and **E**). Identical results were found when defining PCs using in addition surface expression of CXCR4. a chemokine receptor that facilitates homing of PCs to the bone marrow (**Fig. 2D** and **E**; (Hargreaves et al., 2001)). We further investigated the cellular composition of the GFP^low^B220^high^ population that contained the fewest CD138^+^ cells (1 to 20%; **Fig. 2E**). We found that virtually all CD138^neg^ cells within the GFP^low^B220^high^ population expressed the B cell marker CD19 (**Fig. 2F**). Also, and in agreement with *Jchain* gene expression analysis (**Fig. 1D**), these cells were mostly germinal center (GC) B cells (CD38^low^FAS^high^; **Fig. 2F** and **G**). We further performed analyses in the spleen and bone marrow of 12-day SRBC immunized mice heterozygous for the *Jchain*^creERT2^ allele in which precursors and mature B cells, and non-B cell populations were first defined using surface markers and the fraction of GFP expressing cells within those populations determined (**Fig. S1**). We found that most PCs (CD138^+^CXCR4^+^, 60 to 90%) in the spleen and bone marrow expressed GFP (**Fig. S1**). We also observed that a minor fraction of B1b cells (0 to 6%) in the spleen expressed GFP, possibly in agreement with the knowledge that B1b cells are prone to differentiate into PCs and are a source of IgM antibodies during T cell independent responses (Alugupalli et al., 2004). Collectively these data confirmed at the single cell level the gene expression analysis using bulk populations (**Fig. 1D**) and demonstrated that *Jchain* expression is highly enriched in PCs.

**Figure 2.**
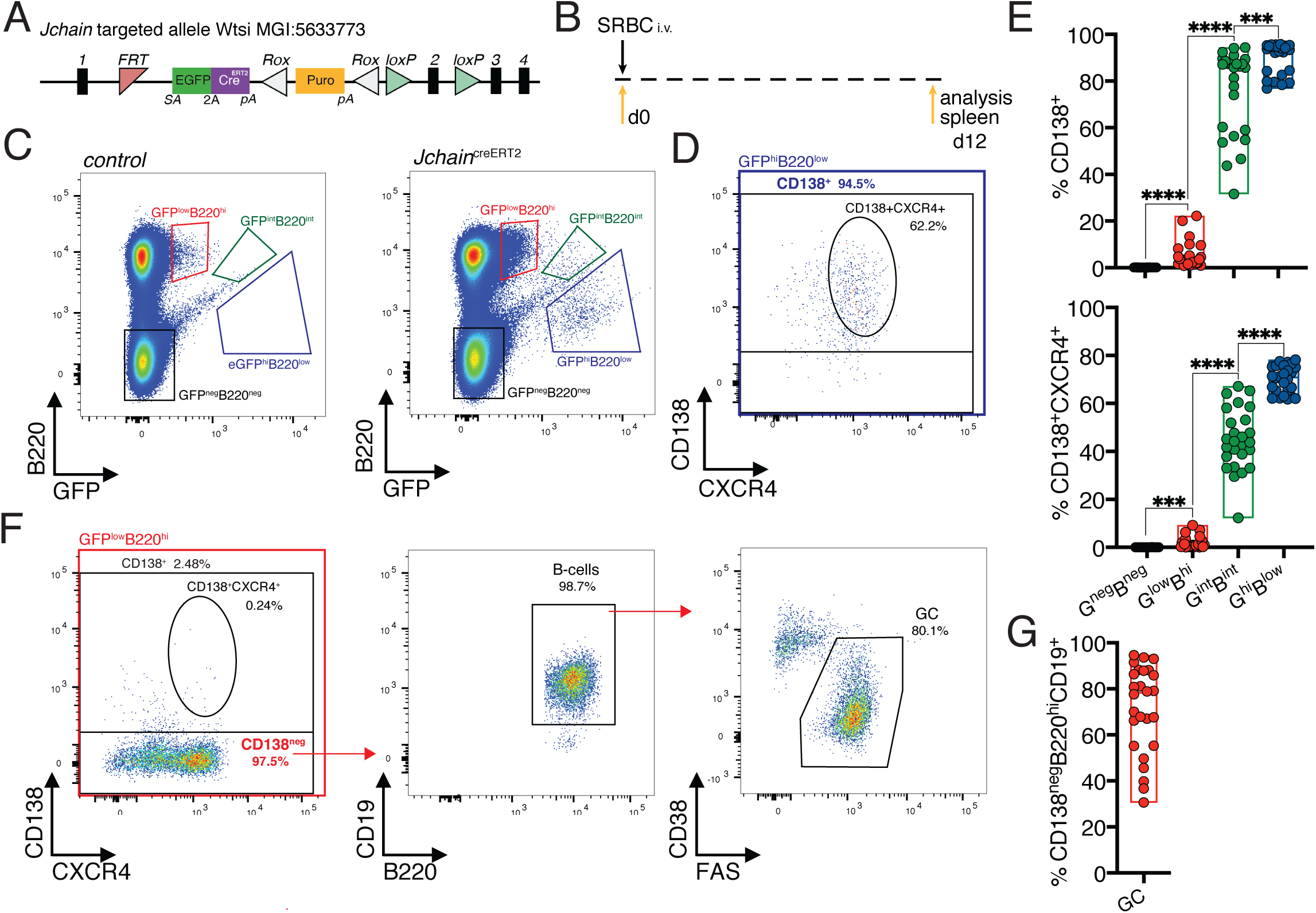
*Jchain* is expressed in a very small fraction of GC B cells and in most Plasma cells. **(A)** Schematic of *Jchain* targeted allele Wtsi MGI:5633773. Rectangular boxes indicate exons, and exon number is on top; pink triangle indicates a FRT sequence; EGFPcre^ERT2^ cassette contains a splice acceptor site (SA)-led EGFP-2A-creERT2 expression cassette followed by a poly-A tail inserted in the intron between exons 1 and 2; white triangle indicates a ROX sequence; orange rectangle indicates a promoter driven puromycin resistance cassette; green triangle indicates loxP sequence. **(B)** Schematic of experimental procedure protocol. Mice carrying the *Jchain*^creERT2^ allele were immunized with sheep red blood cells (SRBC) intravenously (i.v.) on day 0 and spleens of mice were analyzed at day 12 post-immunization. **(C)** Gating strategy of populations by flow-cytometry according to the expression of GFP and B220 in mice carrying the *Jchain*^creERT2^ allele and wild-type B6 mice for negative control of GFP expression. (**D)** Gating strategy by flow-cytometry for plasma cells within the GFP^hi^B220^low^ population using CD138^+^ and CD138^+^CXCR4^+^ markers. **(E)** Cumulative data for CD138^+^ and CD138^+^CXCR4^+^ plasma cells analyzed as in (D). Top: fraction of CD138^+^ plasma cells; Bottom: fraction of CD138^+^CXCR4^+^ plasma cells within the four populations defined by flow-cytometry according to the expression of GFP and B220 in mice carrying the *Jchain*^creERT2^ allele. **(F)** Gating strategy by flow-cytometry for total CD138^+^ and CD138^+^CXCR4^+^ plasma cells within GFP^low^B220^hi^ population. The CD138^neg^ cell fraction within the GFP^low^B220^hi^ population was analyzed for the CD19 B cell marker and stained for CD38 and FAS to determine germinal center (GC) B cells. **(G)** Cumulative data for the frequency of GC B cells within the CD138^neg^GFP^low^B220^hi^ population. Each symbol (E, G) represents an individual mouse; small horizontal lines indicate median, minimum and maximum values. ***=p≤0.001, ****=p≤0.0001 (unpaired Student’s *t*-test). Data are representative of three independent experiments (E, G).

### *Jchain* expression correlates with that of IRF4 and BLIMP1

IRF4 and BLIMP1 transcription factors play an essential role in PC differentiation (Kallies et al., 2007; Klein et al., 2006; Shapiro-Shelef et al., 2003). We analyzed the spleen of 12-day SRBC immunized mice heterozygous for the *Jchain*^creERT2^ allele and determined the expression pattern of IRF4 and BLIMP1 in the populations defined by varied GFP and B220 expression (**Fig. 2C, 3A** and **B**). The GFP^neg^B220^neg^ population was virtually devoid of cells with BLIMP1 and IRF4 expression (**Fig. 3C**). In contrast the GFP^low^B220^high^ cell population contained ∼7% (median) of BLIMP1^+^IRF4^+^ cells (**Fig. 3D** and **E**). These *in vivo* data suggested that *Jchain* expression precedes that of IRF4 and BLIMP1, something previously suggested from *in vitro* cultures of *Blimp1* deficient B cells (Kallies et al., 2007). Still, *Jchain* expression strongly correlated with that of IRF4 and BLIMP1 given that the vast majority of cells within the GFP^int^B220^int^ and GFP^high^B220^low^ populations were BLIMP1^+^IRF4^+^ (**Fig. 3D** and **E**). Overall we identified an *in vivo* population of cells in which *Jchain* expression precedes that of IRF4 and BLIMP1, possibly representing PC precursors. As PC differentiation ensues, *Jchain* expression correlates highly with the expression of the transcription factors BLIMP1 and IRF4 that are critical for the establishment of the PC program.

**Figure 3.**
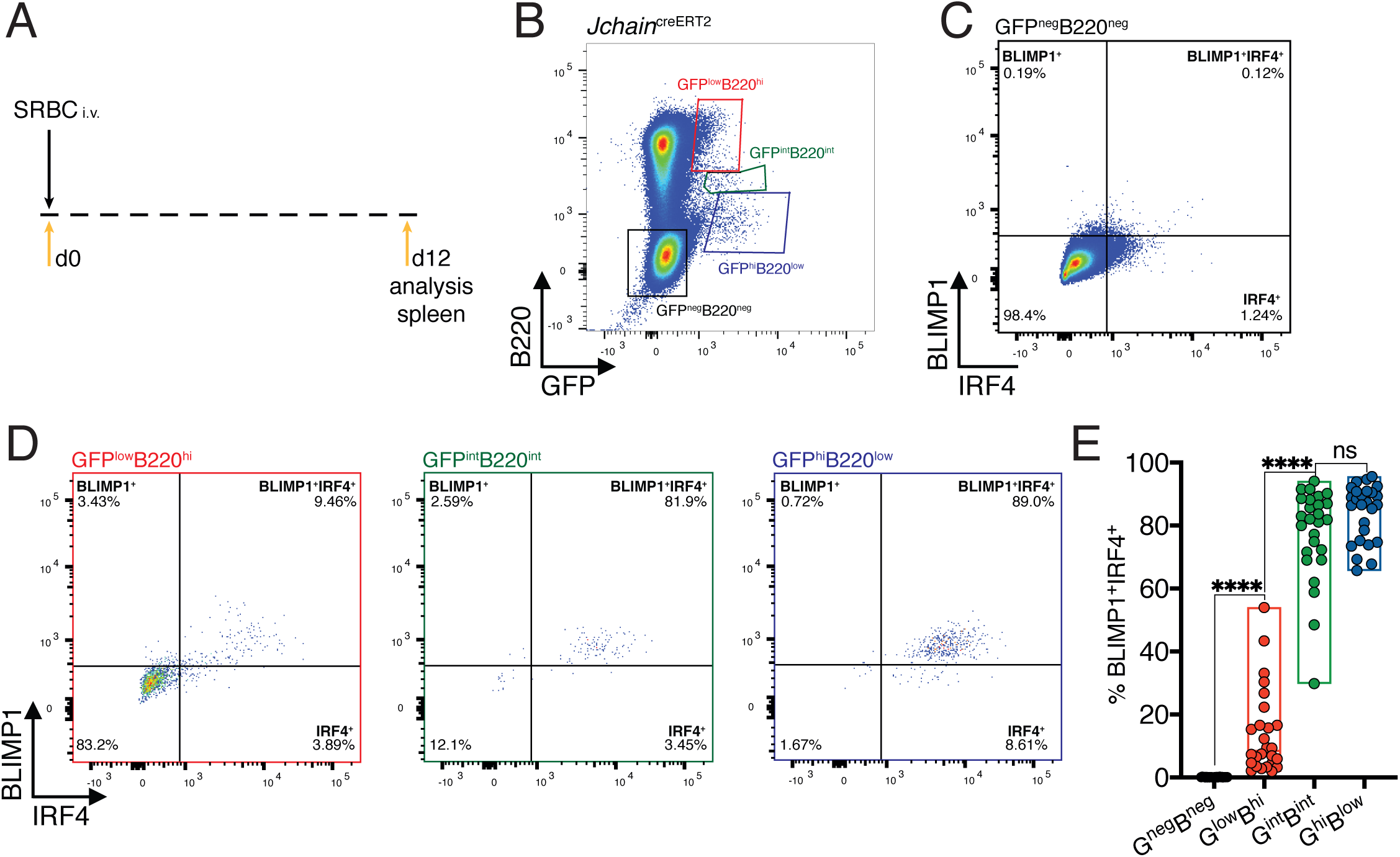
*Jchain* expression correlates with that of IRF4 and BLIMP1. **(A)** Schematic of experimental procedure protocol. Mice carrying the *Jchain*^creERT2^ allele were immunized with sheep red blood cells (SRBC) intravenously (i.v.) on day 0 and spleens of mice were analyzed at day 12 post-immunization. **(B)** Gating strategy of populations by flow-cytometry according to the expression of GFP and B220 in mice carrying the *Jchain*^creERT2^ allele. Wild-type B6 mice for negative control of GFP expression. **(C)** Gating strategy for IRF4 and BLIMP1 expression by flow-cytometry within the GFP^neg^B220^neg^ population. **(D)** Gating strategy for IRF4 and BLIMP1 expression by flow-cytometry within GFP^low^B220^hi^, GFP^int^B220^int^ and GFP^hi^B220^low^ populations, defined as in (B). **(E)** Cumulative data for the frequency of BLIMP1^+^IRF4^+^ cells within the four populations defined as in (B). Each symbol in (E) represents an individual mouse; small horizontal lines indicate median, minimum and maximum values. ns=not significant, ***=p≤0.001, ****=p≤0.0001 (unpaired Student’s *t*-test). Data are representative of three independent experiments (E).

### *Jchain*^creERT2^ mediated genetic manipulation is effective only in plasma cells

We next sought to determine whether the *Jchain*^creERT2^ allele could be used to perform genetic manipulation of PCs. For that we generated compound mutant mice carrying the *Jchain*^creERT2^ allele and a Rosa 26 allele in which RFP expression is conditional to cre-mediated recombination of a loxP-STOP-loxP cassette (*R26*^lslRFP^; (Luche et al., 2007)). In the *Jchain*^creERT2^ allele, cre is fused to an estrogen binding domain (ERT2) that sequesters cre in the cytoplasm through the binding to HSP90 (Feil et al., 2009). Addition of tamoxifen displaces the creERT2-HSP90 complex allowing effective nuclear import of creERT2 and its access to loxP flanked DNA sequences (Feil et al., 2009). We first performed an *in vitro* experiment using a classical plasmablast (B220^low^CD138^+^) differentiation assay in which B cells purified from mice carrying the *Jchain*^creERT2^ and *R26*^lslRFP^ alleles were cultured with LPS in the presence or absence of 4-OH tamoxifen (Andersson et al., 1972). GFP expression was highly enriched in plasmablasts compared to B cells and RFP expression was only observed upon addition of 4-OH tamoxifen to the cell culture, and that occurred virtually only in plasmablasts (**Fig. S2**). This data suggested that creERT2 was specifically expressed by plasmablasts and effectively retained in the cytoplasm in the absence of 4-OH tamoxifen. Similar observations were made in mice carrying the *Jchain*^creERT2^ and *R26*^lslRFP^ alleles *in vivo*. In the absence of tamoxifen administration RFP expressing cells were not detected, confirming the *in vitro* results and supporting that the *Jchain*^creERT2^ allele was not “leaky” in the control of cre activity (**Fig. S1B, C** and **E**). We next immunized mice carrying *Jchain*^creERT2^ and *R26*^lslRFP^ alleles and administered tamoxifen on days 7, 8, 9 and 10 followed by analysis of spleen and bone marrow at day 12 (**Fig. 4A**). Analyses of the populations defined by varied GFP and B220 surface expression (**Fig. 2** and **4B**) revealed that a small fraction of GFP^low^B220^high^ cells were positive for RFP in the spleen (median ∼2%) and in the bone marrow (median ∼1.2%; **Fig. 4C**). In contrast, cre mediated recombination and as consequence RFP expression occurred in ∼37% and ∼31% of GFP^int^B220^int^ in the spleen and bone marrow, respectively, whereas the vast majority of cells within the GFP^high^B220^low^ population had undergone cre mediated recombination and were RFP positive (∼76% in the spleen, and ∼88% in the bone marrow, median; **Fig. 4C** and **D**). These data showed that *Jchain*^creERT2^ mediated cre-recombination was only effective in PC populations validating it as a tool to specifically perform genetic manipulation of PCs.

**Figure 4.**
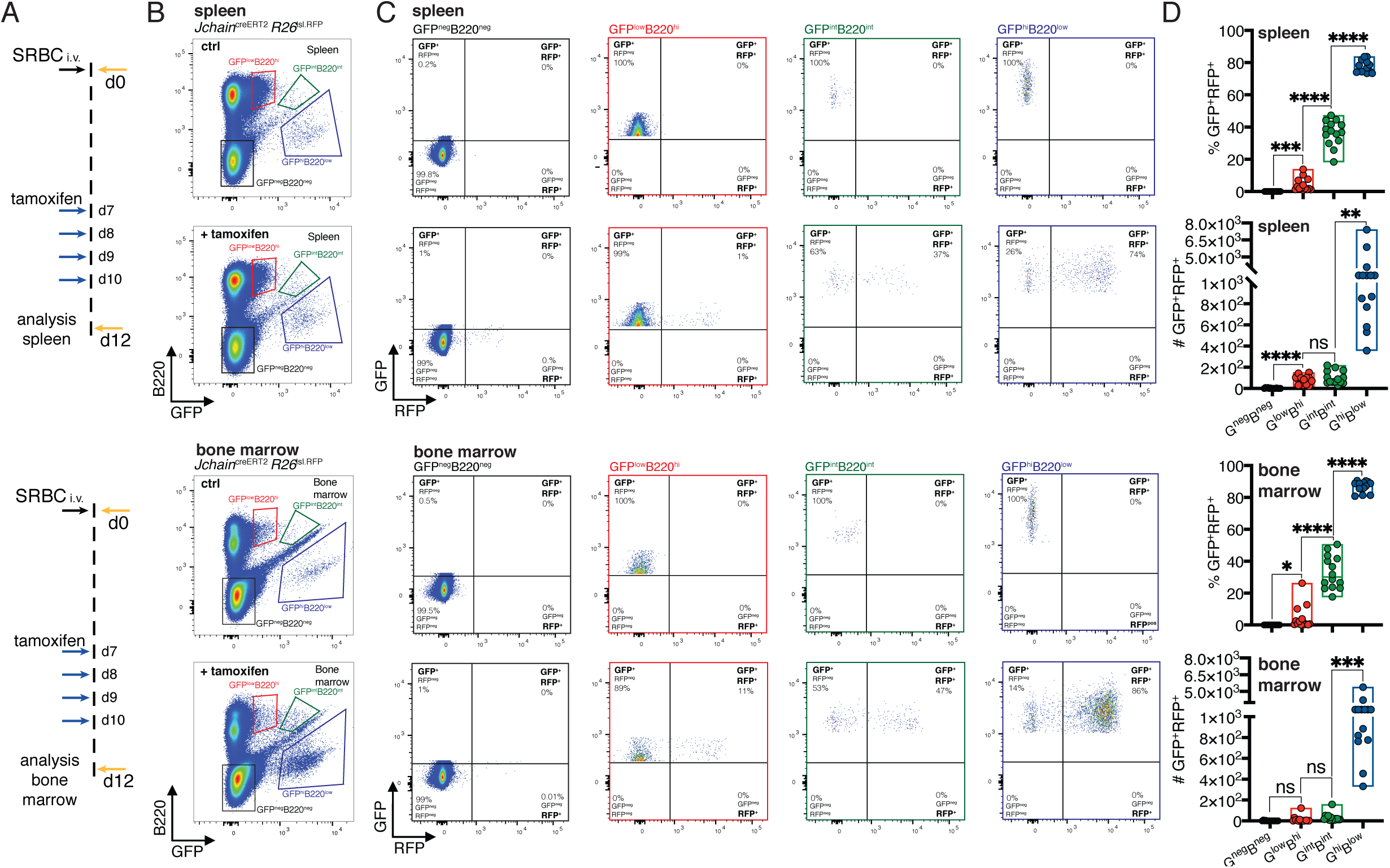
*Jchain*^creERT2^ mediated genetic manipulation is effective only in plasma cells. **(A)** Schematic of experimental procedure protocol. Mice carrying the *Jchain*^creERT2^ and *R26*^lslRFP^ alleles were immunized with sheep red blood cells (SRBC) intravenously (i.v.) on day 0 and spleens (top) and bone marrow (bottom) of mice were analyzed at day 12 post-immunization. A group of mice received tamoxifen treatment for four consecutive days from day 7 to day 10 after immunization. **(B)** Gating strategy of populations by flow-cytometry in spleen (top) and bone marrow (bottom) according to the expression of GFP and B220 in mice carrying the *Jchain*^creERT2^ allele. Wild-type B6 mice for negative control of GFP expression. **(C)** Gating strategy for GFP and RFP expression by flow-cytometry in the four populations defined as in (B). Top: spleen; bottom: bone marrow. **(D)** Cumulative data for the frequency and number of GFP^+^RFP^+^ cells within GFP^neg^B220^neg^, GFP^low^B220^hi^, GFP^int^B220^int^ and GFP^hi^B220^low^ populations, defined as in (B). Top: spleen; bottom: bone marrow. Each symbol (D) represents an individual mouse; small horizontal lines indicate median, minimum and maximum values. ns=not significant, *=p≤0.05, **=p≤0.01, ***=p≤0.001, ****=p≤0.0001 (unpaired Student’s *t*-test). Data are representative of three independent experiments (D).

### Genetic manipulation using *Jchain*^creERT2^ occurs across immunoglobulin isotypes

IgG1 does not multimerize, and due to differences in its secretory tail to that of IgA and IgM, JCHAIN does not associate with IgG1 (Johansen et al., 2000). Currently it is suggested that *Jchain* expression occurs in all PCs regardless of isotype (Castro and Flajnik, 2014; Johansen et al., 2000; Mather et al., 1981). However, this has not been demonstrated at the single cell level. We performed experiments that investigated whether *Jchain*^creERT2^ allele GFP expression (as proxy for *Jchain*), and cre-mediated loxP recombination occurred in PCs across immunoglobulin isotypes. For that we analyzed the spleen, mesenteric lymph nodes (mLN), peyer’s patches and bone marrow of younger (15 weeks) and older (30 weeks) mice carrying the *Jchain*^creERT2^ and *R26*^lslRFP^ alleles. These mice were immunized with SRBC and administered with tamoxifen on days 7, 8, 9 and 10 followed by analysis at day 12 (**Fig. 5A**). We first analyzed total PCs (B220^low^CD138^+^), and within these cells those that expressed GFP (RFP^+^ and RFP^neg^, i.e. *Jchain*^+^) and GFP^+^RFP^+^ cells (i.e. *Jchain*^+^ and cre recombined) to determine the proportions of IgA, IgM and IgG1 expressing cells using intracellular stain (**Fig. 5B-D**). Overall we found only small differences. Analysis of spleens of 15-week-old mice revealed a slight increase in the percentage of IgA^+^ cells within the GFP^+^RFP^+^ PCs only when compared to total PCs (**Fig. 5E**). A similar trend was observed when analyzing the peyer’s patches of 15-week-old mice (**Fig. 5E**). However, in these mice we did not observe differences in the proportion of IgA of other analyzed tissues, nor for IgM and IgG1 in any of the tissues analyzed (**Fig. 5E**). 30-week-old mice showed a slight increase in the percentage of splenic IgM^+^ cells within the GFP^+^RFP^+^ PCs only when compared to total PCs, and for IgA^+^ in the mesenteric lymph node (mLN; **Fig. 5F**). No significant difference were observed in 30-week-old mice for the proportion of IgM or IgA in any of the other analyzed tissues, and in none of the analyzed tissues for IgG1 (**Fig. 5F**). Taken together, these data suggested that *Jchain* expression is not overly represented in IgA or IgM expressing PCs compared to IgG1^+^ PCs. We conclude that *Jchain*^creERT2^ mediated cre-loxP recombination occurs across immunoglobulin isotypes and thus for genetic manipulation also of PCs expressing IgG1.

**Figure 5.**
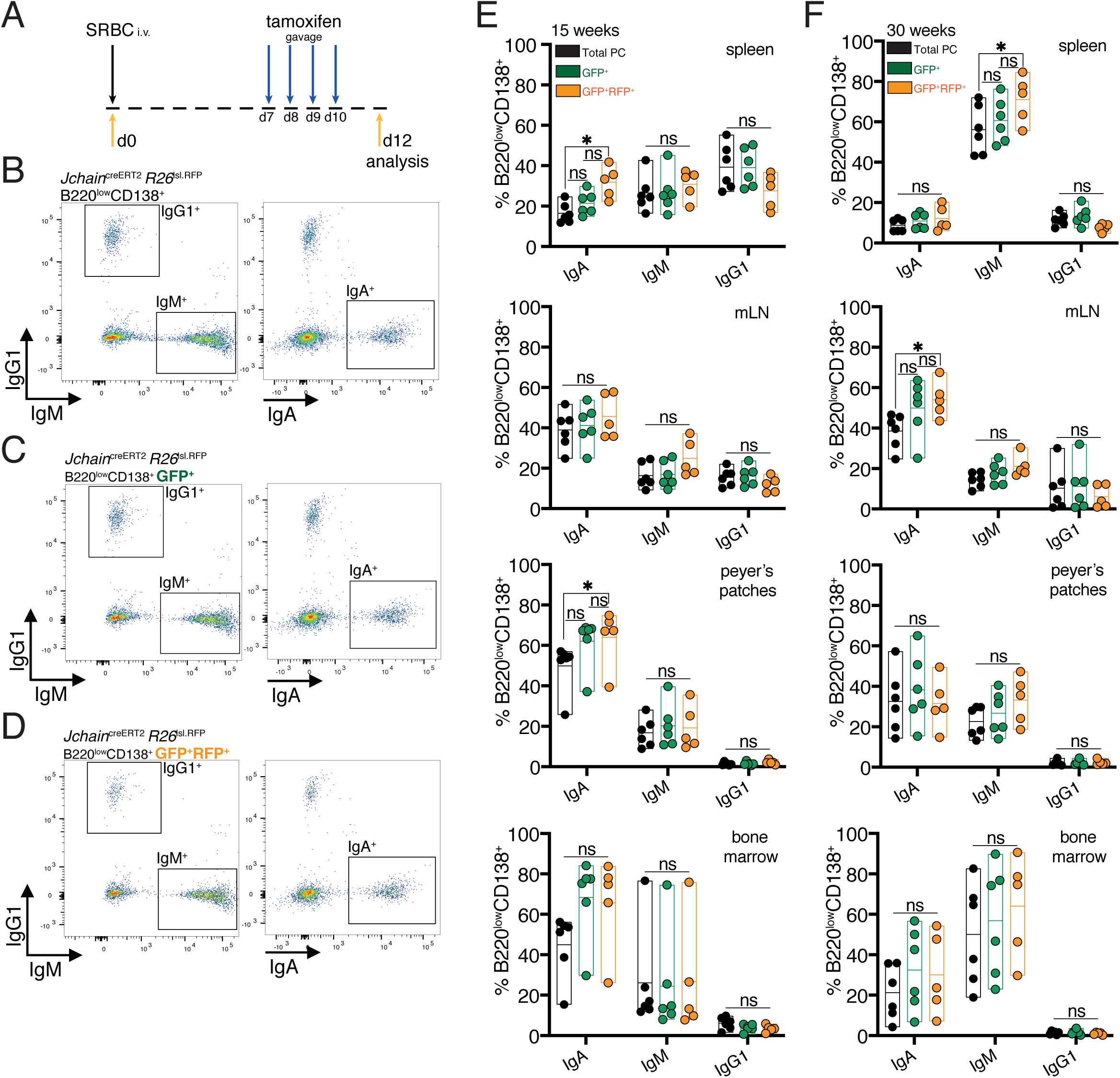
Genetic manipulation using *Jchain*^creERT2^ occurs across immunoglobulin isotypes. **(A)** Schematic of experimental procedure protocol. Mice carrying the *Jchain*^creERT2^ and *R26*^lslRFP^ alleles were immunized with sheep red blood cells (SRBC) intravenously (i.v.) on day 0 and spleens, mesenteric lymph nodes (mLN), peyer’s patches, and bone marrows of mice were analyzed at day 12 post-immunization. Mice received tamoxifen treatment for four consecutive days from day 7 to day 10 after immunization. **(B)** Gating strategy by flow-cytometry for intracellular and extracellular expression of IgG1, IgM and IgA within total B220^low^CD138^+^ plasma cells. Analysis in the spleen is provided as example. **(C)** Gating strategy by flow-cytometry for intracellular and extracellular expression of IgG1, IgM and IgA within B220^low^CD138^+^GFP^+^ (RFP^+^ and RFP^neg^) plasma cells. Analysis in the spleen is provided as example. **(D)** Gating strategy by flow-cytometry for intracellular and extracellular expression of IgG1, IgM and IgA within B220^low^CD138^+^GFP^+^RFP^+^ plasma cells. Analysis in the spleen is provided as example. **(E)** Cumulative data for the fractions of IgA, IgM or IgG1 expressing cells within total plasma cells (PC) (black, B220^low^CD138^+^), GFP^+^ (RFP^+^ and RFP^neg^) plasma cells (green, B220^low^CD138^+^GFP^+^), and GFP^+^RFP^+^ plasma cells (orange, B220^low^CD138^+^GFP^+^RFP^+^) at 15 weeks of age. **(F)** Cumulative data for the fractions of IgA, IgM or IgG1 expressing cells within total plasma cells (PC) (black, B220^low^CD138^+^), GFP^+^ (RFP^+^ and RFP^neg^) plasma cells (green, B220^low^CD138^+^GFP^+^), and GFP^+^RFP^+^ plasma cells (orange, B220^low^CD138^+^GFP^+^RFP^+^) at 30 weeks of age. Each symbol (E, F) represents an individual mouse; small horizontal lines indicate median, minimum and maximum values. ns=not significant, *=p≤0.05 (2way ANOVA). Data are representative of three independent experiments (E, F).

### Inclusive analysis of plasma cell dynamics reveals homeostatic population turnover

Understanding of the PC population turnover is lacking. Multiple investigations have been performed to determine the PC life-span using primarily nucleotide analog DNA incorporation. These studies have provided fundamental insights on PC maintenance and currently it is accepted that a fraction of PCs in the mouse survives for periods longer than 3 months (Ho et al., 1986; Lemke et al., 2016; Manz et al., 1998; Manz et al., 1997; Slifka et al., 1998). However, given the quiescent nature of PCs and that nucleotide analog methodology requires cell division, it is not appropriate to study global population turnover. We investigated the suitability of the *Jchain*^creERT2^ allele to determine the turnover of the PC population in the spleen and bone marrow. We immunized mice carrying the *Jchain*^creERT2^ and *R26*^lslRFP^ alleles at two time-points spaced by a period of 21 days (**Fig. 6A**, (Calado et al., 2010)). Thirty days after the secondary immunization (day 51) we administered tamoxifen for 5 consecutive days to genetically label PCs (**Fig. 6A**). Next we determined the absolute cell number of total PCs, of GFP^+^ cells (RFP^+^ and RFP^neg^, i.e. *Jchain*^+^), GFP^+^RFP^+^ cells (i.e. *Jchain*^+^ cre recombined), and GFP^+^RFP^neg^ cells (i.e. *Jchain*^+^ not cre recombined). We found that over a period of 5 months the cell number of total and GFP^+^ PCs remained constant over time (**Fig. 6B** and **C**). In contrast the cell number of GFP^+^RFP^+^ cells decayed (**Fig. 6B** and **C**) in both spleen (t_1/2_ ∼31.63d) and bone marrow (t_1/2_ ∼251.93d; **Fig. 6D** and **E**). These results agree with the knowledge that the half-life of PCs differs between spleen and the bone marrow (Sze et al., 2000). Notably, analysis of the GFP^+^RFP^neg^ cell numbers revealed that the emergence of these cells paralleled that observed for the decay of GFP^+^RFP^+^ cells (**Fig. 6B** and **C**) both in spleen (t_1/2_ ∼20.20d) and bone marrow (t_1/2_ ∼190.19d; **Fig. 6D** and **E**). These data indicate that the turnover of the *Jchain*^+^ PC population, representing 70 to 90% of all PCs in spleen and bone marrow (**Fig S2B** and **C**), is homeostatically regulated.

**Figure 6.**
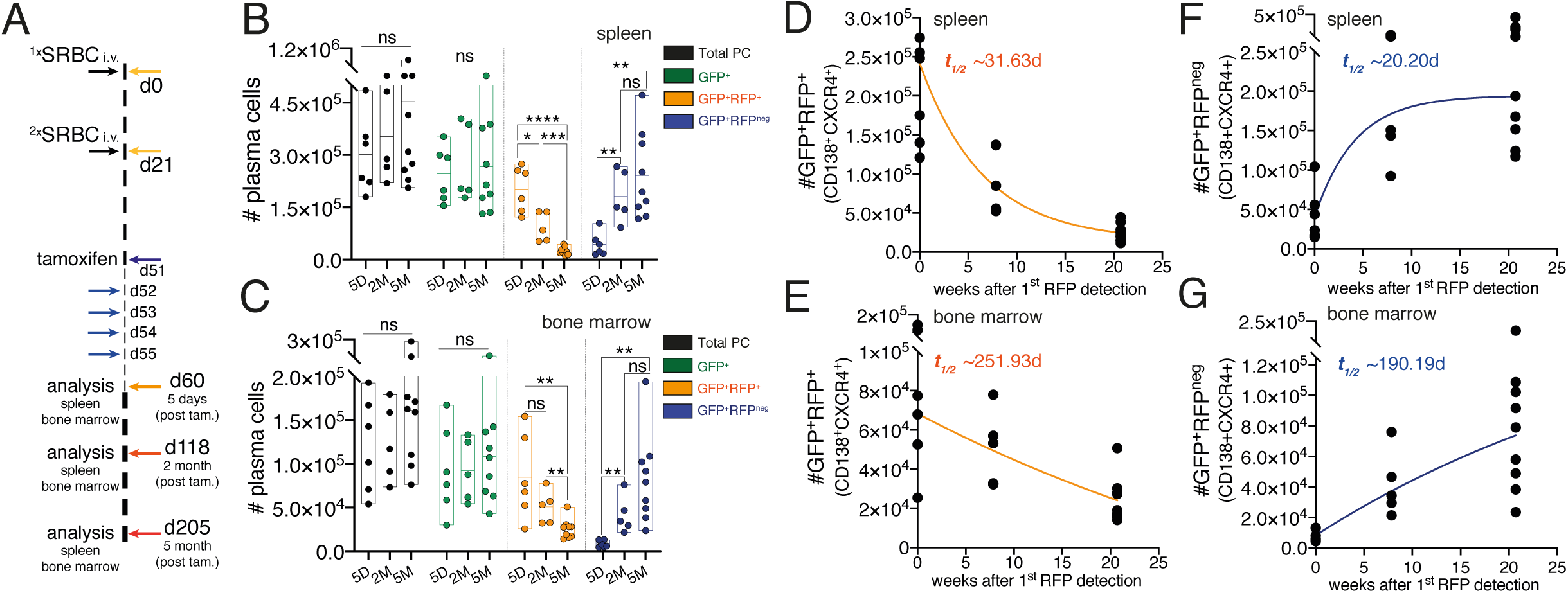
Inclusive analysis of plasma cell dynamics reveals homeostatic population turnover. **(A)** Schematic of experimental procedure protocol. Mice carrying the *Jchain*^creERT2^ and *R26*^lslRFP^ alleles were immunized twice with sheep red blood cells (SRBC) intravenously (i.v.) on day 0 and day 21. Mice received tamoxifen treatment for five consecutive days from day 51 to 55 after the first immunization. Spleens and bone marrow of mice were analyzed at the five-day, two-month and five-month timepoints after the last tamoxifen administration. **(B)** Cumulative data for the absolute cell numbers of total PCs, of GFP^+^ cells (RFP^+^ and RFP^neg^, i.e. *Jchain*^+^), GFP^+^RFP^+^ cells (i.e. *Jchain*^+^ cre recombined), and GFP^+^RFP^neg^ cells (i.e. *Jchain*^+^ not cre recombined) in the spleen. **(C)** Cumulative data for the absolute cell numbers of total PCs, of GFP^+^ cells (RFP^+^ and RFP^neg^, i.e. *Jchain*^+^), GFP^+^RFP^+^ cells (i.e. *Jchain*^+^ cre recombined), and GFP^+^RFP^neg^ cells (i.e. *Jchain*^+^ not cre recombined) in the bone marrow. **(D)** Graphical representation of half-life (t_1/2_) of GFP^+^RFP^+^CD138^+^CXCR4^+^ plasma cells in the spleen using the data presented in (B). Graphing of best-fit curve was performed using the GraphPad Prism 8 software. **(E)** Graphical representation of half-life (t_1/2_) of GFP^+^RFP^+^CD138^+^CXCR4^+^ plasma cells in the bone marrow using the data presented in (C). Graphing of best-fit curve was performed using the GraphPad Prism 8 software. **(F)** Graphical representation of half-life (t_1/2_) of GFP^+^RFP^neg^CD138^+^CXCR4^+^ plasma cells in the spleen using the data presented in (B). Graphing of best-fit curve was performed using the GraphPad Prism 8 software. **(G)** Graphical representation of half-life (t_1/2_) of GFP^+^RFP^neg^CD138^+^CXCR4^+^ plasma cells in the bone marrow using the data presented in (C). Graphing of best-fit curve was performed using the GraphPad Prism 8 software. Each symbol (B-G) represents an individual mouse; small horizontal lines indicate median, minimum and maximum values. ns=not significant, *=p≤0.05 **=p≤0.01, ***=p≤0.001, ****=p≤0.0001 (unpaired Student’s *t*-test). Data are representative of three independent experiments (B, C).

## Discussion

Plasma cells (PC)s are an essential component of the adaptive immune system. However, compared to other B cell lineage populations the molecular analysis of PC gene function *in vivo* has lagged behind. One underlying cause is the lack of tools for specific genetic manipulation in PCs, something available for B cells since 1997, more than 20 years ago (Rickert et al., 1997), and more recently for B cells at multiple stages of development (Boross et al., 2009; Casola et al., 2006; Croker et al., 2004; Crouch et al., 2007; de Boer et al., 2003; Dogan et al., 2009; Duber et al., 2009; Georgiades et al., 2002; Hobeika et al., 2006; Kraus et al., 2004; Kwon et al., 2008; Moriyama et al., 2014; Robbiani et al., 2008; Schweighoffer et al., 2013; Shinnakasu et al., 2016; Weber et al., 2019; Yasuda et al., 2013).

Here we used a systematic analysis of PC associated factors for which expression is increased during PC differentiation (*Xbp1, Jchain, Scd1, Irf4*, and *Prdm1*) with the aim of identifying suitable candidates for the generation of PC-specific tools. *Blimp1*, a critical transcription factor for the establishment of the PC program, displayed an expression pattern unspecific to PCs. This was unsurprising given the knowledge that *Blimp1* is expressed by other hematopoietic and non-hematopoietic cells (John and Garrett-Sinha, 2009), including germ cells (Ohinata et al., 2005). Amongst the analyzed factors we found that *Jchain* displayed the highest RNA expression level in PCs and had the most PC-specific expression pattern (**Fig. 1D**). We considered *Jchain* to be the most suitable candidate for the generation of PC-specific tools.

We searched among alleles generated by the EUCOMMTools consortium and identified a *Jchain*^creERT2^ allele which through a comprehensive series of experiments we validated for specific genetic manipulation of PCs. Briefly, *Jchain* was expressed in a very small subset of B cells (<2%, primarily GC B cells; **Fig. 2F and G;** and **Fig. S1C**) and therefore preceded BLIMP1 and IRF4 expression (**Fig. 3C**). This data agrees with previous analysis of *in vitro* cultures of *Blimp1* deficient B cells in which it was shown that the initiation of *Jchain* expression does not require BLIMP1 (Kallies et al., 2007). Thus the identified population of cells *in vivo* marked by low GFP expression and high surface expression of B220 (GFP^low^B220^hi^), where only ∼7% of cells expressed BLIMP1 (**Fig. 3D** and **E**), likely contains PC precursor cells and future analysis may provide information on the mechanisms underlying the very first steps of PC differentiation.

*Jchain*^creERT2^ mediated genetic manipulation was only effective in PCs (**Fig. 4C** and **D**). In the *Jchain*^creERT2^ allele GFP and cre^ERT2^ are linked by a self-cleaving 2A peptide which ensures a near equitable co-expression of the proteins (Szymczak et al., 2004). As consequence, it is likely that effectiveness of cre recombination correlates with level of cre^ERT2^ because PCs expressed the highest level of *Jchain*, as suggested by the level of GFP expression (**Fig. 4C** and **D**). Of note, the *Jchain*^creERT2^ allele analyzed in this work retained the puromycin selection cassette that is flanked by Rox recombination sites (**Fig. 2A**). This knowledge should be taken in consideration in future experiments involving compound mutant mice that make use of dre recombinase (Anastassiadis et al., 2009). On occasion it has been observed that retention of the selection cassette reduces gene expression of the targeted locus (Meyers et al., 1998; Nagy et al., 1998). As consequence inadvertent or intentional removal of the puromycin selection cassette may potentiate increase creERT2 expression, including in B cells, and as consequence reduce the PC specificity of *Jchain*^creERT2^ mediated genetic manipulation. Lastly, in the *Jchain*^creERT2^ allele the expression of *Jchain* is interrupted by a GFP-2A-creERT2 cassette and induction of cre-mediated recombination deletes *Jchain* exon 2 that is loxP-flanked. It was previously shown that heterozygous deletion of *Jchain* displayed an intermediate phenotype to that of knockout mice (Lycke et al., 1999). This knowledge must be taken into consideration when performing *Jchain*^creERT2^ mediated genetic manipulation, including the need to use the *Jchain*^creERT2^ allele without the targeted manipulation as control, and the utility of the *Jchain*^creERT2^ allele in homozygosity. Future iterations of the *Jchain*^creERT2^ in which JCHAIN protein expression is preserved should be considered.

It is generally assumed that *Jchain* expression occurs in all PCs regardless of their isotypes (Castro and Flajnik, 2014; Johansen et al., 2000; Mather et al., 1981). The characterization of the *Jchain*^creERT2^ allele demonstrated at the single cell level that *Jchain* expression indeed occurs in PCs across immunoglobulin isotypes (**Fig. 5**). However, our analysis did not support the concept that *Jchain* is expressed by all PCs. The occurrence of *Jchain*^+^ and *Jchain*^neg^ is of interest as it may provide insights into PC development and should be a subject of further investigation in the future (Castro and Flajnik, 2014).

Using the *Jchain*^creERT2^ allele we performed inclusive genetic timestamping of PCs, independent of the time at which cells were generated, cell cycle status, and localization (**Fig. 6**). We uncovered that the numbers of total PCs and that of *Jchain*^+^ PCs, which represent 70 to 90% of all PCs (**Fig. 6B** and **C**), remained stable in spleen and bone marrow for at least 5 months (**Fig S2B** and **C**). Notably, within the *Jchain*^+^ PC population, cell decay was compensated by the emergence of new PCs (**Fig. 6B-G**), supporting that PC turnover is homeostatically regulated. Homeostatic control of mature B cell numbers is widely accepted (Crowley et al., 2009). In mature B cells the expression of a B cell receptor is crucial for cell survival (Srinivasan et al., 2009); however, BAFF is the limited resource that defines the boundaries of the biological “space” (Crowley et al., 2009; Srinivasan et al., 2009). On other hand, the regulation of PC turnover remains unclear. PCs express the BCMA receptor that allows the sensing of both BAFF and APRIL, and BCMA deficient mice have reduced PC numbers (O’Connor et al., 2004). However, because BCMA is expressed in B cells committed to PC differentiation further studies are required to disentangle formation and maintenance (Mackay et al., 2003). The *Jchain*^creERT2^ allele is therefore ideally suited to tackle these questions. PC homeostatic regulation could also be the reflection of a limited number of niches in a given organ which would then limit the number of PCs (Hofer et al., 2006; Khodadadi et al., 2019; Lightman et al., 2019; Lindquist et al., 2019; Wilmore and Allman, 2017). Knowledge on the underlying PC turnover mechanisms is important as these are directly related to long-term protection from infection, vaccination and pathogenesis. The *Jchain*^creERT2^ allele is highly suited to for these investigations as it allows genetic manipulation of PCs in vivo in their microenvironment, and to retrieve live time-stamped PCs for downstream analysis.

## Acknowledgments

We thank the members of the Immunity and Cancer laboratory (FCI, London) for critical discussions and review of the manuscript; the FCI BRF and Flow-cytometry platforms for expert advice and technical support.

This work was supported by The Francis Crick Institute which receives its core funding from Cancer Research UK (FC001057), the UK Medical Research Council (FC001057), the Wellcome Trust (FC001057), the CRUK accelerator award EDITOR, and an MRC career development award MR/J008060/1 to D.P.C.

The authors declare no competing financial interests.

## Author contributions

Conceptualization, A.Q.X., R.R.B, D.P.C.; Methodology, A.Q.X., R.R.B, D.P.C.; Investigation, A.Q.X., R.R.B, D.P.C.; Resources, D.P.C.; Writing – Original Draft, A.Q.X., and D.P.C.; Supervision D.P.C.; Funding Acquisition, D.P.C.

## Materials and Methods

### Mice

The *Jchain*^creERT2^ allele was purchased from EMMA in agreement with the Wellcome Trust Sanger Institute (Jchain targeted allele Wtsi MGI:5633773, genebank https://www.i-dcc.org/imits/targ_rep/alleles/43805/escell-clone-genbank-file) and mice were rederived at the The Francis Crick Institute. The allele contains a splice acceptor site (SA), an *EGFP-2A-creERT2* expression cassette and a poly-A tail in the intron between exons 1 and 2 under the *Jchain* promoter. In addition, exon 2 is loxP-flanked and the allele also contains a Rox-flanked puromycin resistance cassette. These mice were crossed to carry a *Rosa26*^lslRFP^ cre recombination reporter allele (*R26*^lsl.RFP^) allele that expresses a non-toxic tandem-dimer red fluorescent protein upon cre-mediated deletion of a floxed STOP cassette (Luche et al., 2007). Mice were maintained on the C57BL/6 background and bred at The Francis Crick Institute biological resources facility under specific pathogen-free conditions. Animal experiments were carried out in accordance with national and institutional guidelines for animal care and were approved by The Francis Crick Institute biological resources facility strategic oversight committee (incorporating the Animal Welfare and Ethical Review Body) and by the Home Office, UK. All animal care and procedures followed guidelines of the UK Home Office according to the Animals (Scientific Procedures) Act 1986 and were approved by Biological Research Facility at the Francis Crick Institute. The age of mice ranged between 15-30 weeks as specified.

### Immunization and *in vivo* induction of cre activity

Mice were injected intravenously with 1 x 10^9^ sheep red blood cells (SRBCs, TCS Biosciences Ltd) in PBS. For the induction of cre activity, 4mg tamoxifen (SIGMA T5648) dissolved in sunflower seed oil were administered by oral gavage to mice once per day for multiple days depending on experimental design.

### *In vitro* B cell culture and induction cre activity

Splenic cells were harvested and B cells were isolated using CD43 (Ly-48) MicroBeads, mouse (Miltenyi Biotec). The purity of B cells was determined by flow-cytometry (>95%). B cells were cultured in 96-well round-bottom plates (Falcon) in B cell media in B cell media (DMEM high glucose/Glutamax from ThermoFisher Scientific supplemented with 10% fetal bovine serum F7524 from Sigma, 100 U/mL Penicillin and 100 µg/mL Streptomycin from Life Technologies, 10 mM HEPES Buffer Solution from Life Technologies, 100 µM MEM Non-essential Amino Acids from ThermoFisher Scientific, 1 mM sodium pyruvate from Life Technologies and 50 uM β-mercaptoethanol from Sigma) at 1 million/mL concentration (200,000 cells/well) with 10ug/mL LPS and varied concentrations of (Z) 4-hydroxytamoxifen (Sigma-Aldrich). Analysis was performed by flow-cytometry at 48, 72 or 96 hours of culture.

### Antibodies

Rat Anti-Mouse Blimp-1, Clone 6D3 (BD Biosciences); Rat Anti-Mouse CD16/CD32, Clone 2.4G2 (BD Biosciences); Rat Anti-Mouse CD19, Clone 1D3 (BD Biosciences); Rat Anti-Mouse CD23, Clone B3B4 (BD Biosciences); Rat Anti-Mouse CD38, Clone 90 (BD Biosciences); Hamster Anti-Mouse CD95, Clone Jo2 (BD Biosciences); Rat Anti-Mouse IgG1, Clone A85-1 (BD Biosciences); Rat Anti-Mouse CD138, Clone 281-2 (BD Biosciences); Rat Anti-Mouse CD45R/B220, Clone RA3-6B2 (BioLegend); Hamster Anti-Mouse CD11c, Clone N418 (BioLegend); Rat Anti-Mouse CD19, Clone 6D5 (BioLegend); Rat Anti-Mouse CD21/35, Clone 7E9 (BioLegend); Rat Anti-Mouse CD43, Clone 1B11 (BioLegend); Rat Anti-Mouse CD86, Clone GL1 (BioLegend); Rat Anti-Mouse BP1, Clone 6C3 (eBioscience); Rat Anti-Mouse CD5, Clone 53-7.3 (eBioscience); Rat Anti-Mouse CXCR4, Clone 2B11 (eBioscience); Rat Anti-Mouse IgA, Clone 11-44-2 (eBioscience); Rat Anti-Mouse IgM, Clone II/41 (eBioscience); Rat Anti-mouse IRF4, Clone 3E4 (eBioscience).

### Flow cytometry

Single cell suspensions were stained with antibodies. We used Zombie NIR Fixable Viability Kit (BioLegend) for live/dead discrimination. For intracellular staining we fixed cells using the BD CytoFix/Cytoperm (BD Biosciences) kit as per manufacturer instructions. Samples were acquired on a BD LSRFortessa analyser using FACSDiva software (BD) and analysed on FlowJo software.

### Quantification and statistical analysis

Data were analyzed with unpaired two-tailed Student’s t test or 2way ANOVA; a p-value of 0.05 or less was considered significant. Prism (v7 and v8, GraphPad) was used for statistical analysis. A single asterisk (*) in the graphs of figures represents a p-value ≤0.05, double asterisks (**) a p-value ≤0.01, triple asterisks (***) a p-value ≤0.001, quadruple asterisks (****) a p-value ≤0.0001, and “ns” stands for not statistically significant, i.e., a p-value >0.05.

### Online supplemental material

Figure S1 displays the analysis of *Jchain* expression across cell lineages in the spleen and bone marrow. These data also demonstrate that *Jchain*^creERT2^ mediated genetic manipulation does not occur in the absence of tamoxifen treatment. Figure S2 shows experiments using a classical LPS-induced B to plasmablast differentiation and the occurrence of *Jchain*^creERT2^ mediated genetic manipulation upon addition of increasing concentrations of 4-hydroxytamoxifen (4-OH-TAM).

## Supplemental Figure Legends

**Figure S1.**
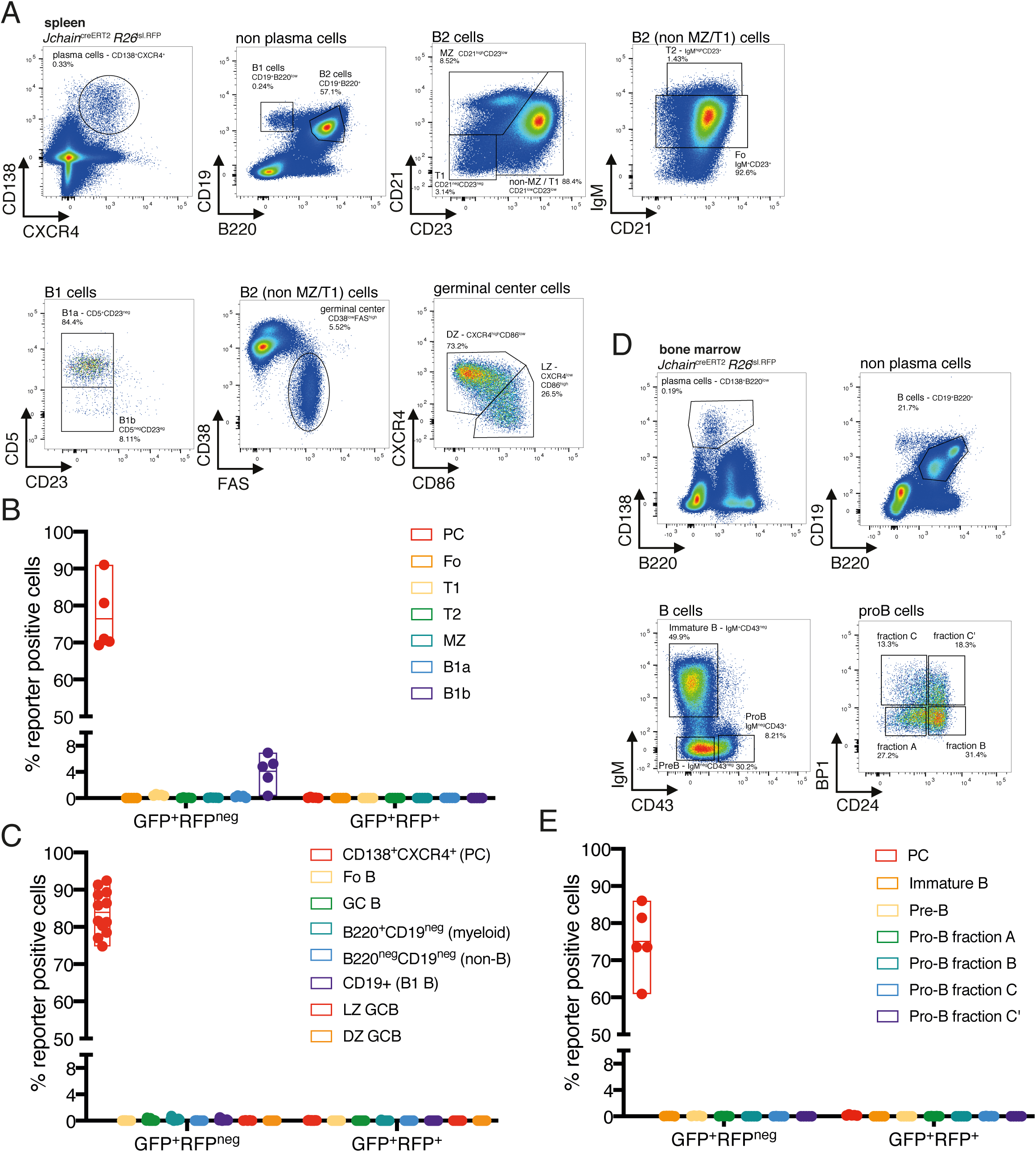
*Jchain* expression in multiple stages of B cell development and other cell lineages. **(A)** Gating strategy by flow-cytometry for the B cell lineage in the spleen. Mice carrying the *Jchain*^creERT2^ and *R26*^lslRFP^ alleles were immunized with sheep red blood cells (SRBC) intravenously (i.v.) at day 0 and spleens were analyzed at day 12. Mice did not receive tamoxifen treatment. Cells were pre-gated on live singlets and for the expression of CD138 and CXCR4 markers. CD138^+^CXCR4^+^ cells were defined as plasma cells (PC). Non-PCs positive for the CD19 and B220 markers were defined as B2 cells, whereas those cells displaying a CD19^+^B220^low^ were defined as B1 cells. B1 cells were further characterized as B1a (CD5^hi^CD23^neg^) and B1b (CD5^low^CD23^neg^) cells. B2 cells were further delineated into marginal zone (MZ) B cells (CD21^hi^CD23^int^), T1 B cells (CD21^low^CD23^low^), T2 B cells (CD21^int^IgM^hi^CD23^hi^) and Follicular B cells (Fo, CD21^int^IgM^int^CD23^hi^. Non MZ or T1 cells were gated for germinal center (GC) B cells (CD38^low^FAS^high^) and these were further delineated into dark zone GC B cells (CXCR4^high^CD86^low^) and light zone GC B cells (CXCR4^low^CD86^high^). (Bonami et al., 2014) **(B)** Cumulative data for fractions of GFP^+^RFP^neg^ and GFP^+^RFP^+^ cells within the B cell lineage in the spleen. Analyzed as in (A). **(C)** Cumulative data for fractions of GFP^+^RFP^neg^ cells and GFP^+^RFP^+^ cells in additional B cell lineage and non-B cell lineage populations in the spleen. Analyzed as in (A). **(D)** Gating strategy by flow-cytometry for the B cell lineage in the bone marrow. Mice carrying the *Jchain*^creERT2^ and *R26*^lslRFP^ alleles were immunized with sheep red blood cells (SRBC) intravenously (i.v.) at day 0 and spleens were analyzed at day 12. Mice did not receive tamoxifen treatment. Cells were pre-gated on live singlets and gated for CD138 and B220 expression. CD138^+^B220^low^ cells were defined as plasma cells (PCs) Non-PCs positive for the CD19 and B220 markers were further examined and further delineated to immature B cells (IgM^+^CD43^neg^), Pre-B cells (fraction D; IgM^neg^CD43^neg^) and Pro-B cells (IgM^neg^CD43^+^). Pro B cells were further sub-divided into Hardy fraction A (BP1^neg^CD24^neg^), fraction B (BP1^neg^CD24^+^), fraction C (BP1^+^CD24^neg^) and fraction C’ (BP1^+^CD24^+^). **(E)** Cumulative data for fractions of GFP^+^RFP^neg^ cells and GFP^+^RFP^+^ cells in the B cell lineage and in plasma cells in the bone marrow. Analyzed as in (D). Each symbol (B, C, E) represents an individual mouse; small horizontal lines indicate median, minimum and maximum values. Data are representative of three independent experiments (B, C, E).

**Figure S2.**
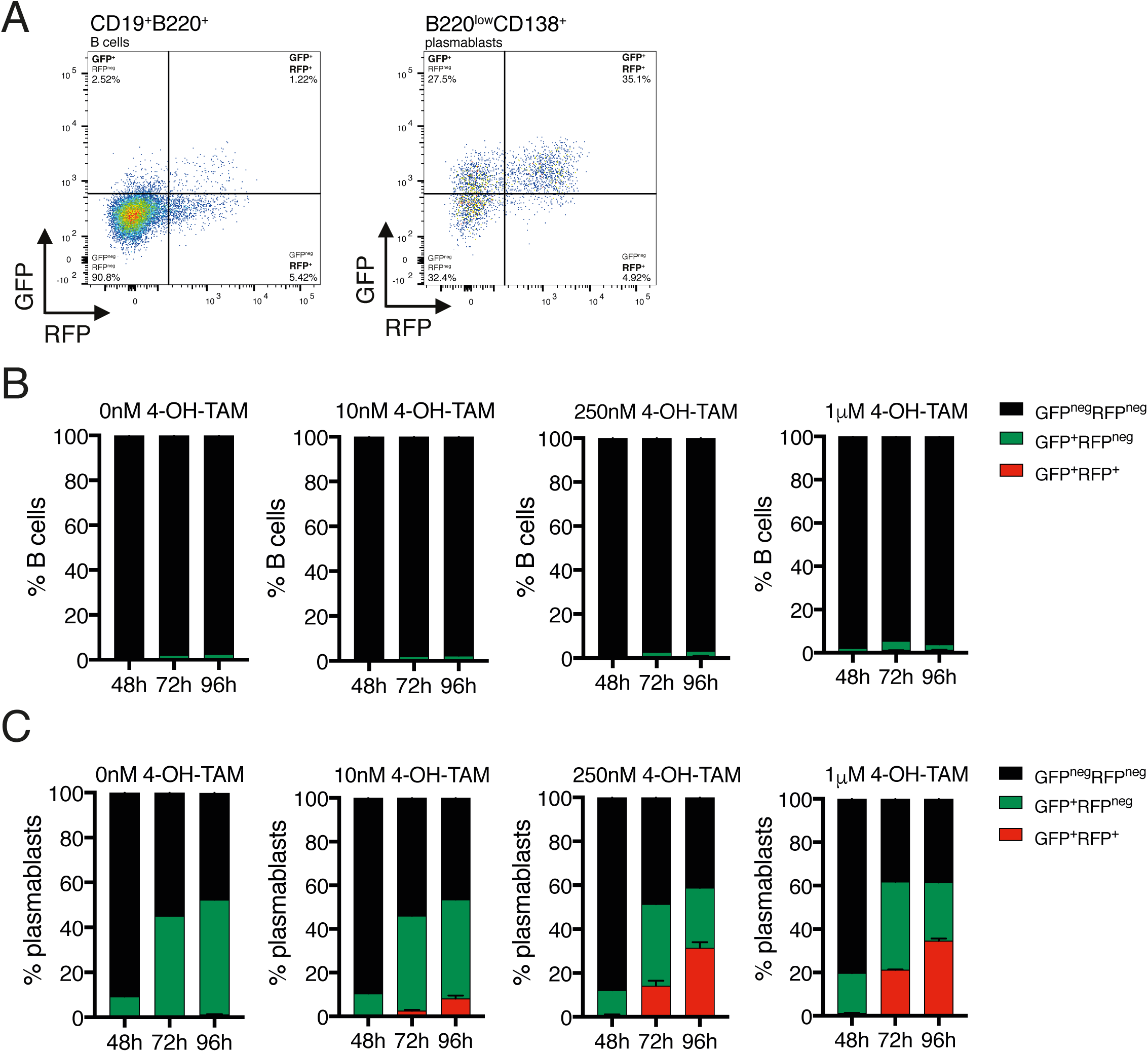
*Jchain*^creERT2^ mediated genetic manipulation in vitro. (**A**) Gating strategy by flow-cytometry for GFP and RFP expression within CD19^+^B220^+^ B cells and B220^low^CD138^+^ plasmablasts. Splenic B cells from unmanipulated mice carrying the *Jchain*^creERT2^ and *R26*^lslRFP^ alleles were enriched using CD43 negative depletion (MACS) and culture *in vitro* in the presence of LPS and 4-hydroxytamoxifen (4-OH-TAM). Cells were analyzed 48h, 72h, and 96h time-points for GFP and RFP expression by flow-cytometry. Data from 96h analysis of a B cell culture with LPS and 1μM 4-OH-TAM is shown as example. (**B**) Cumulative data for fractions of GFP^neg^RFP^neg^, GFP^+^RFP^neg^, GFP^+^RFP^+^ cells within B cells. Analyzed as in (A). (**C**) Cumulative data for fractions of GFP^neg^RFP^neg^, GFP^+^RFP^neg^, GFP^+^RFP^+^ cells within plasmablasts. Analyzed as in (A).

## References

Alugupalli, K.R., J.M. Leong, R.T. Woodland, M. Muramatsu, T. Honjo, and R.M. Gerstein. 2004. B1b lymphocytes confer T cell-independent long-lasting immunity. Immunity 21:379–390.

Anastassiadis, K., J. Fu, C. Patsch, S. Hu, S. Weidlich, K. Duerschke, F. Buchholz, F. Edenhofer, and A.F. Stewart. 2009. Dre recombinase, like Cre, is a highly efficient site-specific recombinase in E. coli, mammalian cells and mice. Dis Model Mech 2:508–515.

Andersson, J., O. Sjoberg, and G. Moller. 1972. Induction of immunoglobulin and antibody synthesis in vitro by lipopolysaccharides. Eur J Immunol 2:349–353.

Bonami, R.H., W.T. Wolfle, J.W. Thomas, and P.L. Kendall. 2014. NFATc2 (NFAT1) assists BCR-mediated anergy in anti-insulin B cells. Mol Immunol 62:321–328.

Boross, P., C. Breukel, P.F. van Loo, J. van der Kaa, J.W. Claassens, H. Bujard, K. Schonig, and J.S. Verbeek. 2009. Highly B lymphocyte-specific tamoxifen inducible transgene expression of CreER T2 by using the LC-1 locus BAC vector. Genesis 47:729–735.

Brandtzaeg, P., and H. Prydz. 1984. Direct evidence for an integrated function of J chain and secretory component in epithelial transport of immunoglobulins. Nature 311:71–73.

Calado, D.P., B. Zhang, L. Srinivasan, Y. Sasaki, J. Seagal, C. Unitt, S. Rodig, J. Kutok, A. Tarakhovsky, M. Schmidt-Supprian, and K. Rajewsky. 2010. Constitutive canonical NF-kappaB activation cooperates with disruption of BLIMP1 in the pathogenesis of activated B cell-like diffuse large cell lymphoma. Cancer Cell 18:580–589.

Casola, S., G. Cattoretti, N. Uyttersprot, S.B. Koralov, J. Seagal, Z. Hao, A. Waisman, A. Egert, D. Ghitza, and K. Rajewsky. 2006. Tracking germinal center B cells expressing germ-line immunoglobulin gamma1 transcripts by conditional gene targeting. Proc Natl Acad Sci U S A 103:7396–7401.

Castro, C.D., and M.F. Flajnik. 2014. Putting J chain back on the map: how might its expression define plasma cell development? J Immunol 193:3248–3255.

Croker, B.A., D. Metcalf, L. Robb, W. Wei, S. Mifsud, L. DiRago, L.A. Cluse, K.D. Sutherland, L. Hartley, E. Williams, J.G. Zhang, D.J. Hilton, N.A. Nicola, W.S. Alexander, and A.W. Roberts. 2004. SOCS3 is a critical physiological negative regulator of G-CSF signaling and emergency granulopoiesis. Immunity 20:153–165.

Crouch, E.E., Z. Li, M. Takizawa, S. Fichtner-Feigl, P. Gourzi, C. Montano, L. Feigenbaum, P. Wilson, S. Janz, F.N. Papavasiliou, and R. Casellas. 2007. Regulation of AID expression in the immune response. J Exp Med 204:1145–1156.

Crowley, J.E., J.E. Stadanlick, J.C. Cambier, and M.P. Cancro. 2009. FcgammaRIIB signals inhibit BLyS signaling and BCR-mediated BLyS receptor up-regulation. Blood 113:1464–1473.

de Boer, J., A. Williams, G. Skavdis, N. Harker, M. Coles, M. Tolaini, T. Norton, K. Williams, K. Roderick, A.J. Potocnik, and D. Kioussis. 2003. Transgenic mice with hematopoietic and lymphoid specific expression of Cre. Eur J Immunol 33:314–325.

Dogan, I., B. Bertocci, V. Vilmont, F. Delbos, J. Megret, S. Storck, C.A. Reynaud, and J.C. Weill. 2009. Multiple layers of B cell memory with different effector functions. Nat Immunol 10:1292–1299.

Duber, S., M. Hafner, M. Krey, S. Lienenklaus, B. Roy, E. Hobeika, M. Reth, T. Buch, A. Waisman, K. Kretschmer, and S. Weiss. 2009. Induction of B-cell development in adult mice reveals the ability of bone marrow to produce B-1a cells. Blood 114:4960–4967.

Feil, S., N. Valtcheva, and R. Feil. 2009. Inducible Cre mice. Methods Mol Biol 530:343–363.

Georgiades, P., S. Ogilvy, H. Duval, D.R. Licence, D.S. Charnock-Jones, S.K. Smith, and C.G. Print. 2002. VavCre transgenic mice: a tool for mutagenesis in hematopoietic and endothelial lineages. Genesis 34:251–256.

Hargreaves, D.C., P.L. Hyman, T.T. Lu, V.N. Ngo, A. Bidgol, G. Suzuki, Y.R. Zou, D.R. Littman, and J.G. Cyster. 2001. A coordinated change in chemokine responsiveness guides plasma cell movements. J Exp Med 194:45–56.

Heng, T.S., M.W. Painter, and C. Immunological Genome Project. 2008. The Immunological Genome Project: networks of gene expression in immune cells. Nat Immunol 9:1091–1094.

Ho, F., J.E. Lortan, I.C. MacLennan, and M. Khan. 1986. Distinct short-lived and long-lived antibody-producing cell populations. Eur J Immunol 16:1297–1301.

Hobeika, E., S. Thiemann, B. Storch, H. Jumaa, P.J. Nielsen, R. Pelanda, and M. Reth. 2006. Testing gene function early in the B cell lineage in mb1-cre mice. Proc Natl Acad Sci U S A 103:13789–13794.

Hofer, T., G. Muehlinghaus, K. Moser, T. Yoshida, E.M. H, K. Hebel, A. Hauser, B. Hoyer, O.L. E, T. Dorner, R.A. Manz, F. Hiepe, and A. Radbruch. 2006. Adaptation of humoral memory. Immunol Rev 211:295–302.

Johansen, F.E., R. Braathen, and P. Brandtzaeg. 2000. Role of J chain in secretory immunoglobulin formation. Scand J Immunol 52:240–248.

John, S.A., and L.A. Garrett-Sinha. 2009. Blimp1: a conserved transcriptional repressor critical for differentiation of many tissues. Exp Cell Res 315:1077–1084.

Kallies, A., J. Hasbold, K. Fairfax, C. Pridans, D. Emslie, B.S. McKenzie, A.M. Lew, L.M. Corcoran, P.D. Hodgkin, D.M. Tarlinton, and S.L. Nutt. 2007. Initiation of plasma-cell differentiation is independent of the transcription factor Blimp-1. Immunity 26:555–566.

Khodadadi, L., Q. Cheng, A. Radbruch, and F. Hiepe. 2019. The Maintenance of Memory Plasma Cells. Front Immunol 10:721.

Klein, U., S. Casola, G. Cattoretti, Q. Shen, M. Lia, T. Mo, T. Ludwig, K. Rajewsky, and R. Dalla-Favera. 2006. Transcription factor IRF4 controls plasma cell differentiation and class-switch recombination. Nat Immunol 7:773–782.

Kraus, M., M.B. Alimzhanov, N. Rajewsky, and K. Rajewsky. 2004. Survival of resting mature B lymphocytes depends on BCR signaling via the Igalpha/beta heterodimer. Cell 117:787–800.

Kwon, K., C. Hutter, Q. Sun, I. Bilic, C. Cobaleda, S. Malin, and M. Busslinger. 2008. Instructive role of the transcription factor E2A in early B lymphopoiesis and germinal center B cell development. Immunity 28:751–762.

Lemke, A., M. Kraft, K. Roth, R. Riedel, D. Lammerding, and A.E. Hauser. 2016. Long-lived plasma cells are generated in mucosal immune responses and contribute to the bone marrow plasma cell pool in mice. Mucosal Immunol 9:83–97.

Lightman, S.M., A. Utley, and K.P. Lee. 2019. Survival of Long-Lived Plasma Cells (LLPC): Piecing Together the Puzzle. Front Immunol 10:965.

Lindquist, R.L., R.A. Niesner, and A.E. Hauser. 2019. In the Right Place, at the Right Time: Spatiotemporal Conditions Determining Plasma Cell Survival and Function. Front Immunol 10:788.

Luche, H., O. Weber, T. Nageswara Rao, C. Blum, and H.J. Fehling. 2007. Faithful activation of an extra-bright red fluorescent protein in “knock-in” Cre-reporter mice ideally suited for lineage tracing studies. Eur J Immunol 37:43–53.

Lycke, N., L. Erlandsson, L. Ekman, K. Schon, and T. Leanderson. 1999. Lack of J chain inhibits the transport of gut IgA and abrogates the development of intestinal antitoxic protection. J Immunol 163:913–919.

Mackay, F., P. Schneider, P. Rennert, and J. Browning. 2003. BAFF AND APRIL: a tutorial on B cell survival. Annu Rev Immunol 21:231–264.

Manz, R.A., M. Lohning, G. Cassese, A. Thiel, and A. Radbruch. 1998. Survival of long-lived plasma cells is independent of antigen. Int Immunol 10:1703–1711.

Manz, R.A., A. Thiel, and A. Radbruch. 1997. Lifetime of plasma cells in the bone marrow. Nature 388:133–134.

Mather, E.L., F.W. Alt, A.L. Bothwell, D. Baltimore, and M.E. Koshland. 1981. Expression of J chain RNA in cell lines representing different stages of B lymphocyte differentiation. Cell 23:369–378.

Max, E.E., and S.J. Korsmeyer. 1985. Human J chain gene. Structure and expression in B lymphoid cells. J Exp Med 161:832–849.

Meyers, E.N., M. Lewandoski, and G.R. Martin. 1998. An Fgf8 mutant allelic series generated by Cre- and Flp-mediated recombination. Nat Genet 18:136–141.

Moriyama, S., N. Takahashi, J.A. Green, S. Hori, M. Kubo, J.G. Cyster, and T. Okada. 2014. Sphingosine-1-phosphate receptor 2 is critical for follicular helper T cell retention in germinal centers. J Exp Med 211:1297–1305.

Nagy, A., C. Moens, E. Ivanyi, J. Pawling, M. Gertsenstein, A.K. Hadjantonakis, M. Pirity, and J. Rossant. 1998. Dissecting the role of N-myc in development using a single targeting vector to generate a series of alleles. Curr Biol 8:661–664.

Nutt, S.L., P.D. Hodgkin, D.M. Tarlinton, and L.M. Corcoran. 2015. The generation of antibody-secreting plasma cells. Nat Rev Immunol 15:160–171.

O’Connor, B.P., V.S. Raman, L.D. Erickson, W.J. Cook, L.K. Weaver, C. Ahonen, L.L. Lin, G.T. Mantchev, R.J. Bram, and R.J. Noelle. 2004. BCMA is essential for the survival of long-lived bone marrow plasma cells. J Exp Med 199:91–98.

Ohinata, Y., B. Payer, D. O’Carroll, K. Ancelin, Y. Ono, M. Sano, S.C. Barton, T. Obukhanych, M. Nussenzweig, A. Tarakhovsky, M. Saitou, and M.A. Surani. 2005. Blimp1 is a critical determinant of the germ cell lineage in mice. Nature 436:207–213.

Palumbo, A., and K. Anderson. 2011. Multiple myeloma. N Engl J Med 364:1046-1060.

Pracht, K., J. Meinzinger, P. Daum, S.R. Schulz, D. Reimer, M. Hauke, E. Roth, D. Mielenz, C. Berek, J. Corte-Real, H.M. Jack, and W. Schuh. 2017. A new staining protocol for detection of murine antibody-secreting plasma cell subsets by flow cytometry. Eur J Immunol 47:1389–1392.

Rickert, R.C., J. Roes, and K. Rajewsky. 1997. B lymphocyte-specific, Cre-mediated mutagenesis in mice. Nucleic Acids Res 25:1317–1318.

Rinkenberger, J.L., J.J. Wallin, K.W. Johnson, and M.E. Koshland. 1996. An interleukin-2 signal relieves BSAP (Pax5)-mediated repression of the immunoglobulin J chain gene. Immunity 5:377–386.

Robbiani, D.F., A. Bothmer, E. Callen, B. Reina-San-Martin, Y. Dorsett, S. Difilippantonio, D.J. Bolland, H.T. Chen, A.E. Corcoran, A. Nussenzweig, and M.C. Nussenzweig. 2008. AID is required for the chromosomal breaks in c-myc that lead to c-myc/IgH translocations. Cell 135:1028–1038.

Schweighoffer, E., L. Vanes, J. Nys, D. Cantrell, S. McCleary, N. Smithers, and V.L. Tybulewicz. 2013. The BAFF receptor transduces survival signals by co-opting the B cell receptor signaling pathway. Immunity 38:475–488.

Shaffer, A.L., M. Shapiro-Shelef, N.N. Iwakoshi, A.H. Lee, S.B. Qian, H. Zhao, X. Yu, L. Yang, B.K. Tan, A. Rosenwald, E.M. Hurt, E. Petroulakis, N. Sonenberg, J.W. Yewdell, K. Calame, L.H. Glimcher, and L.M. Staudt. 2004. XBP1, downstream of Blimp-1, expands the secretory apparatus and other organelles, and increases protein synthesis in plasma cell differentiation. Immunity 21:81–93.

Shapiro-Shelef, M., K.I. Lin, L.J. McHeyzer-Williams, J. Liao, M.G. McHeyzer-Williams, and K. Calame. 2003. Blimp-1 is required for the formation of immunoglobulin secreting plasma cells and pre-plasma memory B cells. Immunity 19:607–620.

Shinnakasu, R., T. Inoue, K. Kometani, S. Moriyama, Y. Adachi, M. Nakayama, Y. Takahashi, H. Fukuyama, T. Okada, and T. Kurosaki. 2016. Regulated selection of germinal-center cells into the memory B cell compartment. Nat Immunol 17:861–869.

Slifka, M.K., R. Antia, J.K. Whitmire, and R. Ahmed. 1998. Humoral immunity due to long-lived plasma cells. Immunity 8:363–372.

Srinivasan, L., Y. Sasaki, D.P. Calado, B. Zhang, J.H. Paik, R.A. DePinho, J.L. Kutok, J.F. Kearney, K.L. Otipoby, and K. Rajewsky. 2009. PI3 kinase signals BCR-dependent mature B cell survival. Cell 139:573–586.

Sze, D.M., K.M. Toellner, C. Garcia de Vinuesa, D.R. Taylor, and I.C. MacLennan. 2000. Intrinsic constraint on plasmablast growth and extrinsic limits of plasma cell survival. J Exp Med 192:813–821.

Szymczak, A.L., C.J. Workman, Y. Wang, K.M. Vignali, S. Dilioglou, E.F. Vanin, and D.A. Vignali. 2004. Correction of multi-gene deficiency in vivo using a single ‘self-cleaving’ 2A peptide-based retroviral vector. Nat Biotechnol 22:589–594.

Weber, T., J. Seagal, W. Winkler, T. Wirtz, V.T. Chu, and K. Rajewsky. 2019. A novel allele for inducible Cre expression in germinal center B cells. Eur J Immunol 49:192–194.

Wilmore, J.R., and D. Allman. 2017. Here, There, and Anywhere? Arguments for and against the Physical Plasma Cell Survival Niche. J Immunol 199:839–845.

Yasuda, T., T. Wirtz, B. Zhang, T. Wunderlich, M. Schmidt-Supprian, T. Sommermann, and K. Rajewsky. 2013. Studying Epstein-Barr virus pathologies and immune surveillance by reconstructing EBV infection in mice. Cold Spring Harb Symp Quant Biol 78:259–263.

